# Conserved and diverged patterns of senescence in *Pristionchus* nematodes

**DOI:** 10.64898/2026.06.26.734768

**Authors:** Rebekah J. White, Cameron J. Weadick

**Affiliations:** Department of Biosciences, Health and Life Sciences, Geoffrey Pope building, University of Exeter, UK; Institute for Neuroscience and Cardiovascular Research, College of Medicine & Veterinary Medicine, Hugh Robson building, George Square, University of Edinburgh, UK

## Abstract

Healthspan, the period of life where organisms are without frailty and/or disease, is a major focus of biogerontological research. To understand late-life decline and increased mortality risk, short-lived organisms such as nematode worms are commonly used. *Pristionchus* nematodes are established models for evolutionary developmental genetics research and show promise as systems for comparative and experimental study of ageing. To support this, we developed phenotypic ageing profiles for the evo-devo model *Pristionchus pacificus* and its little-studied congener *Pristionchus fissidentatus*. We find that various life history traits differ between *P. pacificus* and *P. fissidentatus* (lifespan, brood size, and reproductive period), demonstrating their utility for studying divergent ageing trajectories. Further, several traits are consistently impacted by age, including intestinal barrier function, body size, and locomotory ability. Additionally, in *P. pacificus*, rupture avoidance, cuticle integrity, and feeding rate decline with age, indicating dysregulation across many tissue types. Several age-linked patterns resemble those documented for *Caenorhabditis elegans* despite considerable evolutionary distance, suggesting conserved senescent processes across the Rhabditida family of nematodes. This work highlights similarities and differences in the impact of ageing in two *Pristionchus* nematodes and supports their development as models for evolutionary genetic study of senescence.

## Introduction

Nematodes are the most abundant phylum in the animal kingdom and among the most species-rich invertebrate groups (Bardgett & van der Putten 2014). There is considerable life history variation among nematode species, including alternative reproductive modes, developmental pathways, and ecological lifestyles (Lee 2002, Blaxter 2011), and surveys have revealed striking differences in adult lifespan, with maximum lifespan measured in days, weeks, months, or years, depending on the species (Gems 2000). Much of this lifespan variation reflects differences between long-lived species that are internal parasites of vertebrates, on the one hand, and shorter-lived species that are free-living, on the other. However, notable among-species variation exists even for the short-lived free-living species, which are often much more lab tractable and amenable to controlled comparisons (e.g., Gems 2000, Weadick & Sommer 2016a, Banse et al. 2019, Kern et al. 2023). Understanding the evolutionary and mechanistic bases of lifespan variation is a major aim of biogerontology; short-lived free-living nematodes provide an intriguing system for addressing this problem.

Insights into genetic and environmental influences on lifespan in nematodes mostly derive from study of the free-living worm *Caenorhabditis elegans*, which is now established as a model for studying ageing biology. Research on *C. elegans* has uncovered several genes and pathways that extend life when disrupted, including some with clear human homologs such as AGE-1, DAF-2, and ACN-1 (Friedman & Johnson 1988, Kimura et al. 1997, Kumar et al. 2016, Sutphin et al. 2017), and it is now used as a tool for the anti-ageing drug discovery pipeline (Arya et al. 2010, Ye et al. 2014, Bulterijs & Braeckman 2020, Weinkove & Zavagno 2021). Importantly, much of this work focuses on the ageing process in a general sense, not merely age-dependent survival, for example by exploring age-dependent changes in cell biology, physiology, and behaviour (e.g., Epstein et al. 1972, Bolanowski et al. 1981, Glenn et al. 2004, Herndon et al. 2017). This approach permits consideration of not just lifespan, but healthspan too, and it supports the translation of findings with this model nematode to human biology.

Of course, *C. elegans* is just one nematode, and treating a single species as an exemplar of an entire phylum is limiting: a broader perspective may clarify patterns of conservation and homology that improve translation, and it may reveal novel phenomena in other species that would otherwise be missed. The nematode *Pristionchus pacificus* is a promising system for such work. Like *C. elegans*, *P. pacificus* is a free-living nematode that can be cultured and observed in a laboratory, reared under equivalent conditions (Sommer et al. 1996). Further supporting comparative work, both species have an androdioecious mating system, with the bulk of reproduction happening via hermaphroditic self-fertilisation, and males appearing only rarely under most conditions. The two species have broadly similar life cycles as well, with juveniles hatching from eggs, growing via a series of moults and then maturing within approximately 3-4 days, and then senescing within a few weeks (Byerly et al. 1976, Hong & Sommer 2006).

*P. pacificus* is already established as a “satellite” model system for evolutionary developmental biology within nematodes (Simpson 2002). Early work focused on assessing the generalisability of developmental patterns and processes first explored in *C. elegans* (Félix et al. 1999, Photos et al. 2006, Zauner et al. 2007, Rae et al. 2010, Sinha et al. 2012a, b). However, later work revealed that *Pristionchus* nematodes possess developmental and behavioural features not seen in *C. elegans* or its close relatives, such as a plastic mouthform polyphenism and associated behavioural differences that facilitate a predatory diet, making it biologically interesting in an independent sense (Bento et al. 2010, Moczek et al. 2011, Serobyan et al. 2014, Werner et al. 2017, Sieriebriennikov et al. 2018, Namdeo et al. 2018, Sommer 2020). Efforts to study these features in *P. pacificus* led to the development of genomic resources and tools for genetic manipulation, including RNAi mediated gene knockdown, CRISPR/Cas9 mediated knock-out, embryo transgenesis, and single worm transcriptomics (Pires da Silva 2006, Cinkornpumin & Hong 2011, Witte et al. 2014, Namai & Sugimoto 2018, Sun et al. 2021), as well as the collection and phylogenetic study of many wild strains and closely related species (Morgan et al. 2012, Rödelsperger et al. 2014). Combined, these tools and resources make *P. pacificus* well suited for both mechanistic and evolutionary study of ageing biology.

A few studies have explored aspects of lifespan and senescence in *P. pacificus* and its close relatives, and collectively they demonstrate the suitability of this system for ageing biology research. For instance, use of *P. pacificus* has shown the connection between lifespan and innate immunity in germline-ablated individuals, which live longer and have increased resistance to bacterial pathogens, indicating a role for somatic germline signalling in immunity-linked senescence (Hsin & Kenyon 1999, Patel et al. 2002, Rae et al. 2012). Comparative studies, conducted under standard rearing conditions, revealed considerable variation in longevity among *P. pacificus* strains and different *Pristionchus* species, thus demonstrating natural genetic variation in senescence pattern (Weadick & Sommer 2016a, 2017). Additionally, like *C. elegans*, *P. pacificus* exhibits death fluorescence (Coburn & Gems 2013) and semelparous reproductive death (Kern et al. 2020).

To aid the development of *P. pacificus* into a model system for comparative and mechanistic investigation of ageing biology, we set out to describe healthspan indicators in *P. pacificus* with the goal of determining which tissues can be assayed to observe changes with age. Our aim was to evaluate a variety of known/suspected hallmarks of ageing and determine whether the senescent patterns seen in *C. elegans* occur similarly or differently in *P. pacificus*. We complemented this with study of another self-fertile androdioecious *Pristionchus* species, *P. fissidentatus* (Kanzaki et al. 2012). Within the scope of the *Pristionchus* phylogeny, *P. pacificus* and *P. fissidentatus* are distant relatives, but compared to *C. elegans* they are close relatives (Blaxter 2011). By including this additional species in our study, we aimed to further clarify how conserved ageing processes are in nematodes, and provide a foundation for in depth study of ageing across the *Pristionchus* genus. Using a combination of time-to-event analyses and young-vs-old comparison experiments, we found quantifiable age-dependent changes in life history, behaviour, and tissue integrity in *P. pacificus*. For selected traits we confirmed similar patterns in *P. fissidentatus* but also identified distinct patterns of late-life mortality risk in the two species. Combined, this work provides new insights into the nature of senescence in *Pristionchus* nematodes and furthers the development of *P. pacificus* as a model for mechanistic and evolutionary ageing research.

## Methods

### NEMATODE STRAINS AND CULTURE CONDITIONS

The nematode strains used were *P. pacificus* RS2333 and *P. fissidentatus* RS5133. Strain maintenance followed standard protocols (Stiernagle 2006). In brief, nematodes were raised at 20 ± 1°C in 6.0 cm diameter petri dishes on standard nematode growth medium (NGM) and fed an ad lib diet of *Escherichia coli* OP50. *E. coli* OP50 was prepared from single colonies grown at 37 °C for ca. 20 h in low-salt LB media, with agitation, then refrigerating the liquid culture prior to use; NGM plates were seeded with 300–400 µL of *E. coli* OP50 that was then allowed to form a lawn before usage. Because both strains are self-fertile, with life cycles of ∼3–4 days, worms were transferred to fresh plates every 2-3 days to avoid overcrowding and starvation. Worms were transferred to new plates using a heat-sterilised platinum-iridium wire pick (individual worms) or by washing plates with M9+ buffer (bulk collections). No efforts were made to enrich for male nematodes: all assays were conducted using unmated hermaphrodite worms. Neither antibiotics nor antifungals were included in the NGM or the food source. The worms were cultured under these conditions for several generations prior to the start of experiments. The nematode and bacterial strains were obtained from the Sommer Lab at the Max Planck Institute for Biology Tübingen.

Age-synchronised populations of worms were obtained by bleaching (Stiernagle 2006). In brief, eggs and gravid hermaphrodites were washed from plates with M9+ buffer, incubated in a caustic bleach solution for 8 minutes with periodic vortexing, then washed twice and incubated overnight at room temperature (ca. 19–21°C) in M9+ without food. J2-stage juveniles that hatched overnight were collected via centrifugation at 1300 rcf, washed again in M9+, concentrated, and transferred to OP50-spotted NGM plates and incubated at 20 ± 1°C. These worms would reach the J4 pre-adult stage ∼2 days later, at which point hermaphrodites were collected for use in lifespan assays or maintained (with periodic transfers) until of appropriate age for comparative young-vs-old assays; in such cases, we compared young (J4+3 day) versus old (J4+12 day) adult hermaphrodites, as justified below.

### TIME TO EVENT ANALYSES

Lifespan/healthspan assays of unmated hermaphrodite worms were conducted using conditions similar to those described above for general worm culturing: worms were fed ad lib *E. coli* OP50 and maintained on NGM agar at 20 ± 1°C. Replicate assay plates (3.5 cm diameter petri dishes spotted with 50 µL OP50) were loaded with 10 J4-stage worms per plate, then observed daily until all worms had died. Worms were transferred to new plates every 2-3 days during the reproductive period (hence, 2-3 transfers), but thereafter only transferred if required due to deteriorating plate conditions (e.g. contamination) to minimise further handling. Worms suspected as dead were gently prodded with a platinum work pick, with lack of response taken as evidence of worm death; observations were taken using a Zeiss Stemi 508 microscope outfitted with 10X oculars and 0.63X-5.0X zoom. The following events were recorded: (1) death with no clear cause, (2) death due to vulval rupturing, (3) death due to “bagging” (i.e. internal hatching of progeny), and (4) death due to escape (i.e. desiccation on the plate walls). Rupturing was characterised as ejection of internal organs from the vulva, as described in (Leiser et al. 2016). Bagging was characterised as hatched juvenile(s) present in the body of an adult (Herndon et al. 2003). Time spans are reported in days since hatching (*n* days). The final sample size for *P. pacificus* was n = 940 worms (from 94 plates distributed across 19 independent experiments) and n = 350 for *P. fissidentatus* (from 35 plates across 8 independent experiments).

For survival analysis, Kaplan-Meier (KM) survival curves were used to summarise and visualise survival patterns within each of the two species, and Cox Proportional-Hazards (Cox PH) models used to compare survival patterns between the species (Kaplan & Meier 1958, Cox 1972, Cox & Oakes 1984). Deaths due to bagging, or escapes were treated as censored data points when analysing longevity, but additional analyses were done treating bagging and rupturing as biological events that are potential markers of age-dependent worm health (Mosser et al. 2011, Leiser et al. 2016). The proportionality assumption underlying the Cox PH model was checked with a *χ*2 test and graphical examination of the scaled Schoenfeld residuals (Schoenfeld 1982, Hess 1995). Extreme outliers were detected (and removed) by examining deviance residuals. Survival analyses were conducted using the following packages for R (v4.3.0): survival (v3.2.11), survminer (v0.4.9), and ggfortify (v0.4.17) (Therneau 2015, Tang et al. 2016, R Core Team 2019).

### BROOD SIZE

For the brood size assays, J4-stage worms were individually housed on 3.5 cm plates spotted with 50 µL *E. coli* OP50 and moved to new plates every day until egg laying stopped. Old plates were maintained for 2-3 days, at which point hatched larvae were counted. Worms that died or escaped before Age_fin_ (latest age of reproduction) were omitted from calculations of brood size and reproductive period. Reproductive period is defined as the time between first and final days where the average number of offspring, plus standard deviation (sd), is more than 1. The sample size for this assay was >30 for both species, with measurements collected across four independent experiments.

### BODY SIZE

To determine body size, worms were placed on 2 % agarose pad slides and imaged with an Olympus SZX16 microscope with SFD PLAPO 1XPF lens at 11.5x zoom magnification, using cellSens camera control software (v.4.2.1). Images of worms were straightened to permit measurement of body length (from the tip of the mouth to the end of the body cavity) and width (measured above perpendicular to the body length at the vulva) using Worm-align macro within FIJI v1.53f (Schindelin et al. 2012, Okkenhaug et al. 2020).. At least 10 worms per age across three replicate experiments were included. A two-way analysis of variance (ANOVA), followed by a post-hoc Tukey’s honestly significant difference (HSD) Test, was used to examine the effects of age and species on body size.

### CUTICLE PERMEABILITY

A protocol for cuticle permeability to acridine orange (AO) was adapted from previously described methods (Xiong et al. 2017, Van de Walle et al. 2019). Worms were washed off plates, stained with 5 μg/mL acridine orange in M9+ buffer for 15 minutes with gentle agitation, followed by three M9+ washes (with worms collected via centrifugation at 1300 rcf), then mounted on 2 % agarose pads and imaged using an Axiophot Fluorescence Microscope (Zeiss) and the tetramethylrhodamine (TRITC) filter set. AO accumulation was imaged with an exposure setting of 50% target. Fluorescence intensity was measured using FIJI v1.53f (Schindelin et al. 2012) and converted to an Intensity score (IS) that was calculated as *IS_worm_ = IntDen_worm_ - (Area_worm_×F_background_)*, where IntDen is integrated density and F is fluorescence (adapted from (“Measuring cell fluorescence using ImageJ — The Open Lab Book v1.0,” n. d.). Data was then analysed with a t-test to test for a difference in fluorescence between young adult and old worms.

### INTESTINAL BARRIER FUNCTION

To assess intestinal barrier function, worms were suspended in a solution containing both food and a dye, then examined for introgression of ingested dye particles through the intestinal wall into the pseudocoelom (Gelino et al. 2016). The food-dye mixture was prepared by dissolving erioglaucine disodium salt in standard *E. coli* OP50 liquid culture (final concentration of dye = 0.05 g/mL). Batches of worms were incubated in 100 µL food-dye mixture for 150 minutes at room temperature, washed 3-4 times, as described above, until the blue dye was no longer apparent, and then moved to 2 % agarose pads for imaging at 6.3x and 11.5x magnification with an Olympus SZX16 microscope with SFD PLAPO 1XPF lens. Dye can be visible in the intestine, pharynx, ovaries, and pseudocoelom. Worms were scored on a binary yes/no system of dye present in the pseudocoelom, while ignoring dye present elsewhere to ensure it permeated through the cuticle rather than entered via the mouth, anus or vulva, then the percentage of individuals scored as having dye introgression in each batch was calculated. For each age and species, three independent experiments were carried out, with at least 13 worms per age across three replicates. A two-way weighted ANOVA was used to test for a difference between ages and species, with each batch’s percentage weighted by the number of worms assayed. Colour contrast was enhanced to equivalent degrees in young vs. old images.

### MOVEMENT TRACKING

To track worm movement, small batches of worms (between 10-35 washed worms per replicate assay) were moved to spotted NGM plates and left to acclimatise for 20 minutes at room temperature. The NGM plates had been spotted within the last 24h to ensure a thin bacterial lawn and minimal variation across plates. A 10 second video was recorded at 65 frames per second and x6.3 magnification with a VWR Visicam TC 20 PLUS camera.

Several body size and locomotory parameters were then measured with WormLab software (MBF Bioscience, Williston, VT; figure 3B) (Okkenhaug et al. 2020) across the 650 frames of video from each assay: worm length, worm width, wavelength, mean amplitude, track length, smoothed speed, and turn count. Any individual that could not be outlined by the software was removed from the study (for example being obscured by a thicker edge of the OP50 lawn or crawling directly above a plate label). This was repeated three times, with data deriving from 35-50 worms per age for each species. Metrics were defined per the WormLab manual.

Analysis of the locomotory data was conducted using R (v4.3.0) (R Core Team 2019). For track length and smoothed speed, some frame values represented reversals and were negative, so all were converted into absolute values. We then performed a principal component analysis (PCA) to explore differences between ages and identify associations between locomotory variables and body size measurements, using package ggbiplot (v0.6.2) and PCAtest (v0.0.1) (Camargo 2022, Vu 2024). The effect of body size parameters and age (young vs old) on the locomotory metrics were tested via multivariate analysis of variance (MANOVA). Outliers were identified with the identify_outliers function from the rstatix (v0.7.2) (Kassambara 2023) package and outliers outside the 95 % ± 1.5 IQR were removed. Normality assumption of the residuals was checked with a qq plot and Shapiro-Wilks test.

### FEEDING RATE

To test for feeding rate differences between young and old worms, we examined ingestion of fluorescent microspheres, using a protocol modified from (Fueser et al. 2020). *E. coli* concentration was determined by measuring optical density at 600 nm using a SpectraMax M5 (Molecular Devices) spectrophotometer and converting OD600 to cells/mL using a conversion factor of 8x10^8^ (Myers et al. 2013). In brief, 0.125 µl of Fluoresbrite® YG Microspheres (Polysciences,18860-1) was mixed with 50 µl of 10^8^ cells/ml OP50, then spotted onto a 3.5 cm NGM plate and air-dried overnight. Worms were left to crawl on unspotted NGM plates for 30 minutes to dislodge old OP50, then transferred to a microsphere-OP50 spotted plate to feed for 20 minutes. Worms were then washed and heat-killed by incubating at 80 °C for 5 minutes, before mounting on 2 % agarose pad on a glass slide. Slides were imaged using an Axio Observer Z1 (Zeiss) microscope with 4′,6-diamidino-2-phenylindole (DAPI) filter set, and beads were manually counted. This was only studied in *P. pacificus*, with 24 worms per age group distributed across four replicates. Data was analysed in R with a one-way ANOVA, with the mean count weighted by the number of contributing worms, to determine if there was a difference in number of beads ingested by young and old adults.

## Results

### TIME-TO-EVENT ANALYSIS

There was a moderate but statistically significant difference between lifespans of the two *Pristionchus* species (figure 1A-B). *P. pacificus* had a longer lifespan with a median of 23 days (95 % CI 23-25) with 26.1 % of starting individuals censored. The latest age which had a survival of more than 10 % was day 34. This is consistent with (Gilarte et al. 2015) where the average lifespan was 22.5 days (± 7.3 sd), but shorter than (Weadick & Sommer 2016) which had a median of 37 days (95 % CI 33–39) since maturation (around 35 days since hatching; note this study used a slightly different NGM recipe (including tryptone in place of peptone) and incubated worms at a slightly cooler temperature (19 °C rather than 20 °C), and therefore delayed development and life cycle is expected). *P. fissidentatus* had a median lifespan of 21 (19-22) days with 47.6 % censored individuals (more than half of censors were due to bag-of-worms within the first 8 days; supplemental table 1). The latest age which had more than 10 % of worms remaining (including censored individuals) was day 26.

**Figure 1:**
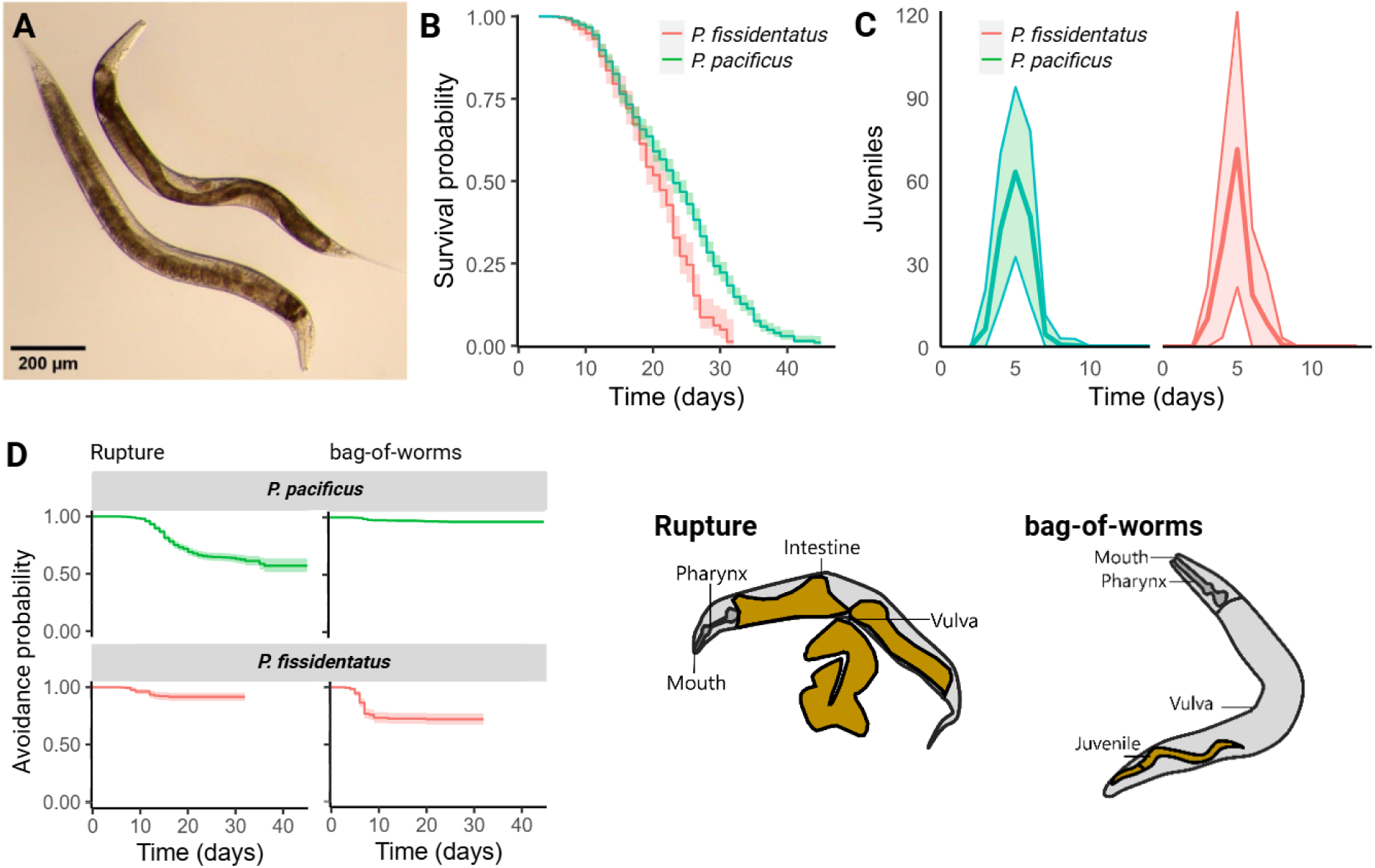
Lifespan, brood size, rupturing probability, and bag-of-worms probability of *Pristionchus* nematodes. A. *P. pacificus* (right) and *P. fissidentatus* (left) young adults (J4+3 days since hatching) raised at 20 °C and suspended in M9+ buffer. B. Survival curves of *P. pacificus* and *P. fissidentatus*. Ribbon = 0.95 confidence interval. C. Average number of juveniles per day in each species. C. Lifetime bag-of-worms and rupturing phenotypes for worms at 20 °C. Left: KM curves of rupture and bag avoidance frequency. Ribbon = 0.95 confidence interval. Right: Diagram of an adult nematode exhibiting rupture (top) and bagging (bottom), with the distinguishing feature of each phenotype highlighted in yellow.

*P. pacificus* ruptured more than *P. fissidentatus*, and rupturing events occurred in an age-dependent manner. In *P. pacificus*, 29.0 % of individuals ruptured. More than half of these events fell within days 12 to 22, with proportions of ruptured individuals increasing with age up until day 26 where the plot plateaued (figure 1D). The risk window continued until 37 days. In *P. fissidentatus*, 12.8 % ruptured, and the proportion of ruptured individuals increased with age up until day 13. There was a significant difference in rupture avoidance probability between species, with *P. pacificus* rupturing more (Cox PH p < 0.05). Notably, in both species, rupture data did not meet the proportional hazards regression assumption meaning the hazard ratio changed over time (chisq = 23.2, df = 1, p = 1.4e-06).

In *P. pacificus,* 3.1 % individuals bagged, whereas in *P. fissidentatus*, 26.6 % bagged. The KM plots indicate that the risk window for bagging is largely restricted to the self-fertile reproductive period for both species. The risk window for bagging in *P. fissidentatus* was between 4 and 9 days, after which risk plateaued (Figure 1D).

### REPRODUCTIVE PERIOD AND BROOD SIZE

*P. pacificus* produced an average of 164 selfed juveniles per individual (± 8.83 sd; table 1; figure 1C). All selfed offspring were produced within seven days of the mother’s maturation. The reproductive peak occurred on day 5 (Age_max_), where an average of 63 progeny (Juv_max_) were produced per mother. *P. fissidentatus* produced an average of 147 juveniles per individual (± 11.67 sd), and all selfed offspring within six days of maturation. Juv_max_ was 71, with Age_max_ also being on the fifth day.

### BODY SIZE AND LOCOMOTION

Body size significantly increased with age in both species, particularly for body width (Figure 2). In *P. pacificus*, width increased from 87 μm (± 7.1 sd) to 125 μm (± 11.8 sd) when comparing young (J4+3 days) and old (J4+ 12 days) adult worms, an increase of 44%. Meanwhile, length increased by 27%, from 790 μm (± 58.7 sd) to 1002 μm (± 92.4 sd). Similar results were seen in *P. fissidentatus*, where width increased by 32%, growing from 118 μm (± 15.5 sd) to 156 μm (± 21.6 sd), and length increased by 11%, from 954 μm (± 122.0 sd) to 1065 μm (± 92.4 sd). *P. fissidentatus* worms were larger, overall, being significantly wider and longer than *P. pacificus* worms when considering young adults, and significantly wider for old adults. These results suggest broadly but not completely similar body size growth dynamics in the two *Pristionchus* species.

**Figure 2:**
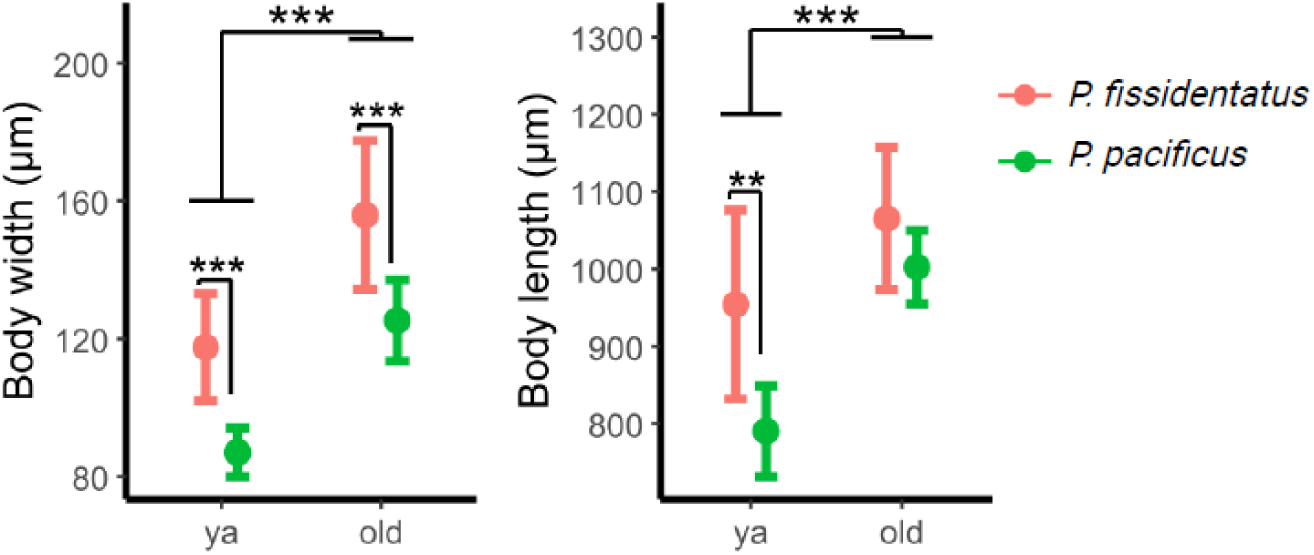
Body size measures (width and length) of young adult (3 days post maturation) and old (12 days post maturation) *Pristionchus* nematodes. *** is p > 0.001, ** is p > 0.01, no asterisk is ns. Bars = sd, n = 12.

**Figure 3:**
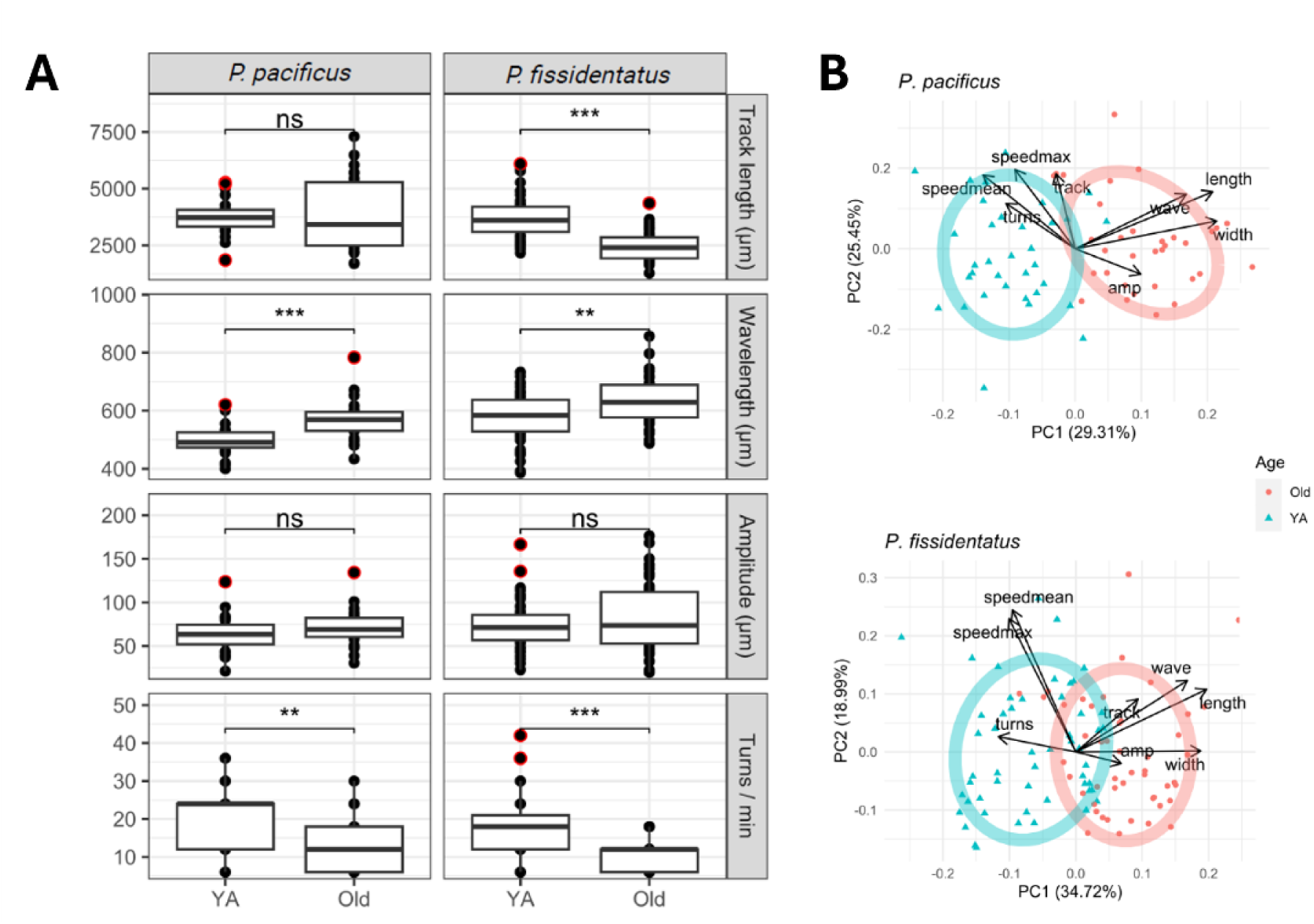
Locomotion metrics comparing young and older adults. **A.** Box plots of track length, wavelength, and amplitude and turn counts of each species. Each point is an individual worm average. Red circled point = outlier datum. For asterisks, *** = p < 0.001, ** = p < 0.01, ns = not significantly different. **B.** PCAs of locomotory and body size metrics in young and old worms. Width = body width, length = body length, wave = wavelength of worm movement, track = distance travelled. Ellipses representing confidence interval level set at 68 % (representing approximately 1 standard deviation of the mean), indicating PC1 is due to age. Note lack of strict correlation between track length and speeds, which is due to track being measured by centre point versus speed being measured by head position. In this standardised PCA, the angles between variable vectors approximate the correlations among the variables.

Multivariate analysis of locomotion parameters, measured in young and old worms, revealed age-linked changes in movement in both species. First, significant differences in locomotory parameters were revealed by MANOVA tests for both species (*P. pacificus*: F(1, 44) = 22.219, p < 0.001; *P. fissidentatus*: F(1, 87) = 22.220, p = < 0.001). Second, PCAs of locomotion and body size measurements uncovered qualitatively similar age-dependent patterns in the two species. For both species, permutation tests revealed significant correlational structure and two significant PC axes (all p < 0.05), with PC1 and PC2 accounting for 29.3 % and 25.5 % of the variance, respectively, for *P. pacificus*, and 34.7 % and 19.0 %, respectively, for *P. fissidentatus*. Variation along PC1 was linked to age group (particularly for *P. pacificus*, where age-group separation along PC1 was stronger): turn rates and crawling speed were increased in young adults, whereas wavelength, amplitude, and (as expected) body size were higher in old worms, suggesting age-dependent declines in activity and flexibility. Track length was not clearly linked with age in *P. pacificus* but was increased in old worms in *P. fissidentatus*. Variation along PC2 may represent naturally-occurring individual-level variation in crawling behaviour and/or the contribution of plate-level batch effects.

We then considered the individual parameters within species (supplemental table 2). In *P. pacificus*, on average old adults made 12 fewer turns per minute (changes of direction) than old adults, while old *P. fissidentatus* made 6 fewer turns. Likewise, in both species, there was a significant difference in wavelength (worm ‘bendiness’) between the age groups. In old worms, this increased by 70.68 µm in *P. pacificus*, and by 59.13 µm *P. fissidentatus*. The proportional wavelength increases with age resembles the proportional increase seen in body size with age. Track length also increased in old worms by 1246 µm in *P. fissidentatus*, but not in *P. pacificus*. The mean speed also declined with age in both species, with old *P. pacificus* slower by 38.11 µm/s and *P. fissidentatus* by 26.09 µm/s.

### INTESTINAL BARRIER FUNCTION

To assess intestinal barrier function, worms were suspended in an *E. coli* and blue dye mix and observed for dye leakage through the intestine and into the pseudocoelom. Overall, there was a significant difference in leakage between young and old adults (F(1, 2) = 13.41, p = 0.006) indicating that intestinal barrier function declined with age. The magnitude of the young vs old difference did not differ between species (F(1, 2) = 1.08, p = 0.320), indicating they show a similar leakage and decline with age. In *P. pacificus*, blue dye in the young adult worms predominantly remained in the intestine and sometimes ovaries (likely through the vulva). Across three batches, blue dye accumulated in parts of the pseudocoelom in 6 % of the 34 young adults and 34 % of the 41 old worms, however the extent of dye introgression varied greatly among individuals (figure 5A-B). Of the 77 young adult *P. fissidentatus*, 17 % had blue dye in the pseudocoelom, compared with 35 % of the 48 older worms. Older adults had a build-up of enlarged unfertilised oocytes (figure 5C).

**Figure 4:**
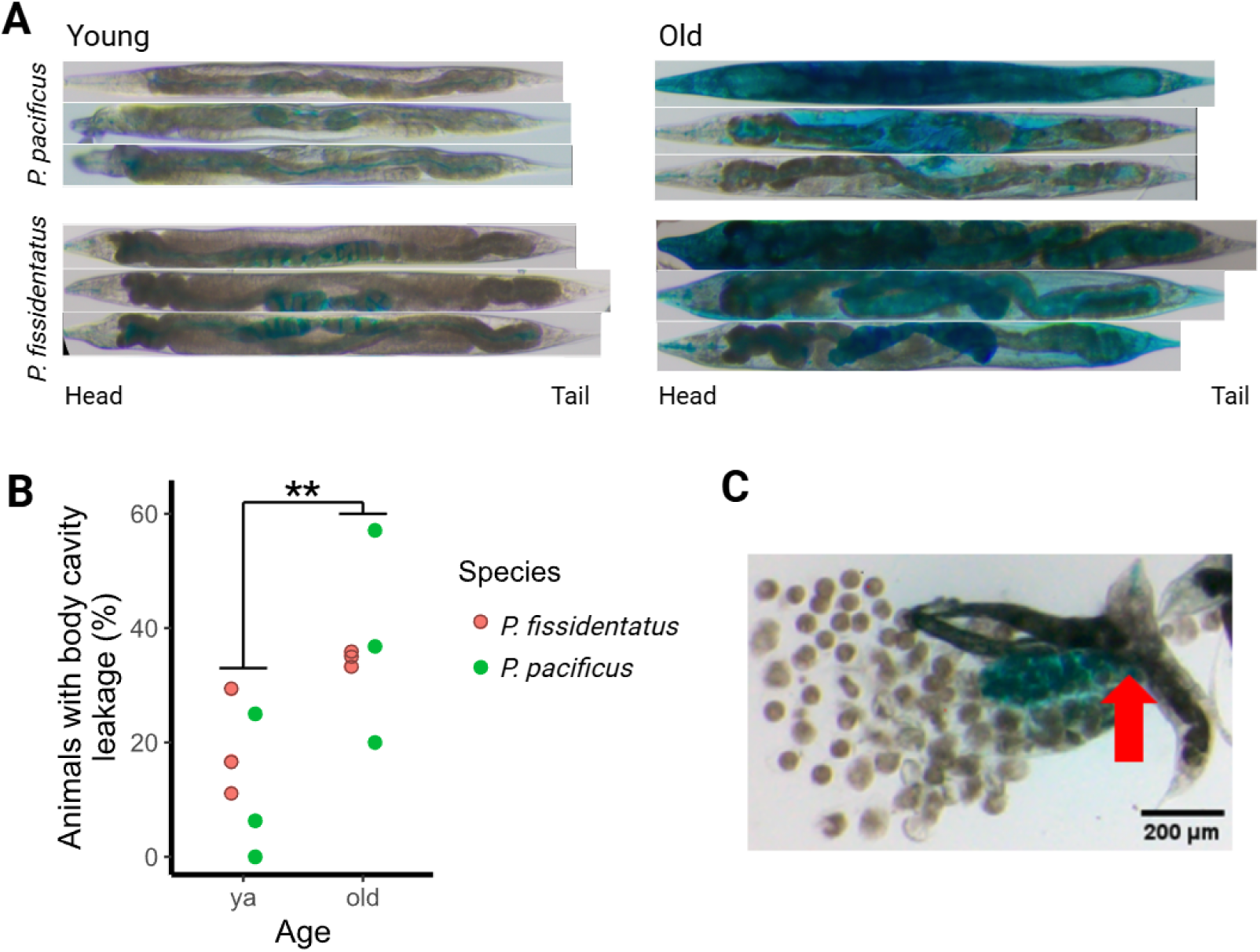
Intestinal barrier function assay. A. Young worms accumulate dye in the intestine, pharynx, ovary. By contrast, older adults also accumulate dye in the pseudocoelom (body cavity) in some individuals. *P. fissidentatus* show less transparent bodies due to a combination of concentrated dye in pseudocoelom and build-up of unfertilised oocytes. B. Intestinal barrier function of *Pristionchus* nematodes in young and old adults, showing mean percentage of animals with pseudocoelom leakage. One point = one batch of 3-36 worms, ** = p > 0.01 (weighted ANOVA to account for varying batch sizes. *P. pacificus* YA = 34, old = 41, and *P. fissidentatus* YA = 77, old = 48). C. An old *P. fissidentatus* where the body is ruptured and shrunken and contains enlarged oocytes. Blue dye is still present in the ovaries external to the body. Red arrow = rupture site, vulva.

**Figure 5:**
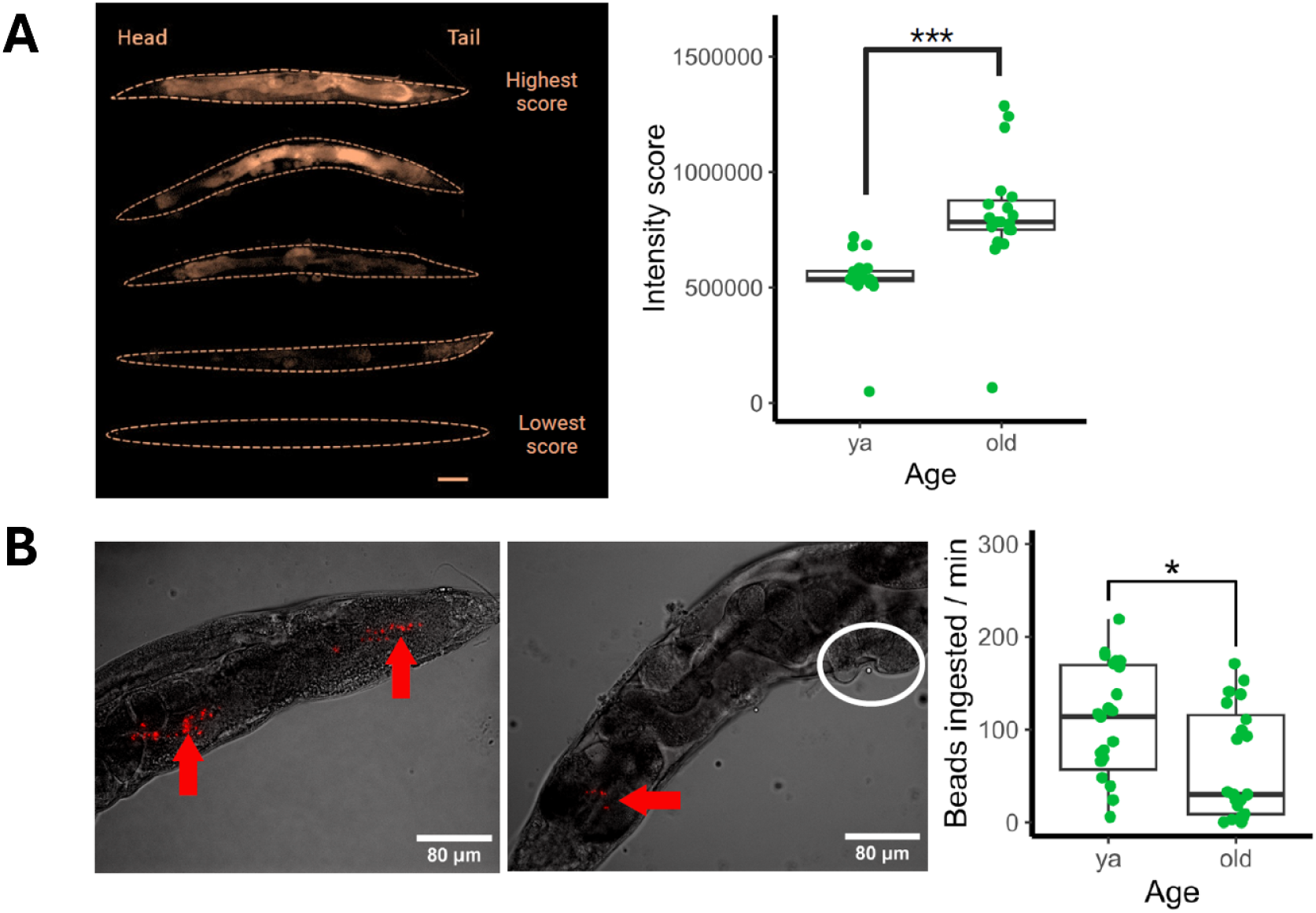
Cuticle permeability and feeding rate of young adult and old *P. pacificus* hermaphrodites. A. AO absorption after 15 minutes. Left: Examples of worms with varying intensity scores, scale bar = 100 µm. Right: Average intensity score of young adult (J4+3 days) and old (J4+12 days) worms, n = 20 per age group. B. Feeding rate after 15 minutes on solid NGM seeded with OP50-microsphere mix and heat-killed. Left: Young adult tail. Middle: Old adult below pharynx. Red arrows = bead accumulation, circle = blister-like effect of heat-killing on the cuticle. Right: Average bead ingestion rate, n = 24 per age group.

### CUTICLE INTEGRITY AND FEEDING RATE

Cuticle integrity was measured by soaking young and old worms in a fluorescent nucleotide dye and exposing them to ultraviolet light to observe how much moved through the cuticle to stain the internal tissues. *P. pacificus* showed an increase in intensity score with age F(1, 38) = 20.14, p < 0.001), increasing by 56.9 % from 575106 AU (± 125217 sd) to 902596 AU (± 293656 sd; figure 6A). This indicates that more AO dye was able to pass through the cuticle and into the body. The intestine was typically stained more than other features. This may have been due to ingestion differences with age, so this was assessed in *P. pacificus* by allowing young and old worms to feed on OP50 containing fluorescent beads, then counting the number of beads in the intestine. Beads accumulated throughout the intestine, often just past the pharynx or before the anus. Young worms ingested an average of 113 beads (± 81.1 sd) during the 20 minute incubation period, which was significantly more than old worms, which ingested just 62 beads (± 59.2 sd; figure 6B) despite being larger animals (F(1, 23) = 6.32, p = 0.016), indicating that feeding rate decreased. This suggests that the increased AO staining reported above is not due to an increase in ingestion.

## Discussion

We characterised patterns of ageing in two self-fertile species of *Pristionchus* nematodes, *P. pacificus* and *P. fissidentatus*. The two species have similar lifespans under laboratory conditions and therefore provide a useful contrast for examining patterns of senescence beyond survivorship. By noting patterns of age-dependent decline due to rupturing and bagging, and by testing for differences between young and old worms for a variety of behavioural and physiological traits, we see that ageing impacts the two species similarly in some ways, but differently in others. Firstly, age-dependent risk patterns for vulval rupturing or matricide differ markedly between the two: ageing *P. pacificus* hermaphrodites are much more likely to die of vulval rupturing than matricidal (internal) hatching of progeny, whereas the opposite is seen in *P. fissidentatus*. Secondly, for both species we see that older worms are bigger, slower, and have reduced intestinal integrity. Additionally, we found that cuticular integrity and feeding rate both decline with age in *P. pacificus*. Below, we discuss these findings in light of past work on ageing in *C. elegans* and *Pristionchus* nematodes.

Rupturing and bagging are severely damaging health events that rapidly lead to death in nematodes, including *C. elegans* (Luc et al. 1979, Chen & Caswell-Chen 2003, Leiser et al. 2016). Both events are plausibly linked to declining reproductive health: rupturing occurs when internal organs are expelled through the vulva, whereas bagging occurs when delayed egg laying leads to internal hatching of progeny. Under this viewpoint, measuring the onset of rupturing and bagging in ageing populations of worms is a simple way to gauge senescent reproductive health (Pickett & Kornfeld 2013, Leiser et al. 2016). Despite this, rupturing and bagging events are often censored during data analysis for survival assays (Lionaki & Tavernarakis 2013) and are only rarely examined as traits of interest. Our observation of different risk patterns for rupturing and bagging in two *Pristionchus* species (reared and observed in a common environment) indicates that, despite their similar life cycles and lifespans, reproductive health declines with age differently in *P. pacificus* and *P. fissidentatus*. This finding supports efforts to ensure bagging and rupturing outcomes are reported in nematode survival studies (e.g. Zhao et al. 2017), and our work provides a foundation for future work exploring whether interventions can impact these easy-to-identify senescent patterns in *Pristionchus* nematodes.

The nematode body maintains high internal hydrostatic pressure (Harris & Crofton 1957, Park et al. 2007), and rupturing could result from weakened ability to tolerate normal levels of internal pressure, abnormally increased internal pressure, or both. *P. pacificus* worms were particularly susceptible to rupturing, with the rate of rupturing rising considerably as the worms aged out of the self-fertile reproductive period, and then declining gradually thereafter. This could indicate that rupturing is a risk associated with dysregulated egg-laying behaviour, and/or the accumulation of vulval wear-and-tear damage associated with producing over 100 progeny within the first few days of adulthood. Leiser et al. 2016 speculated that an increased risk of rupturing in post-reproductive *C. elegans* hermaphrodites was due to elevated internal pressure caused by accumulation of bacterial food in the gut, and showed that reducing food intake lowered rupturing risk, as did disrupting the key insulin-type receptor gene *daf-2*. Our worms were fed *ad lib*, and so were not experimentally deprived of calories, but we found that older *P. pacificus* hermaphrodites have reduced food intake rate, which could contribute to the reduced rupturing risk seen in very advanced ages. As such, one possible explanation for our observation of reduced rupturing risk at very advanced ages is that reduced food intake rates results in reduced internal pressure on the vulva. Pressure on the vulva may also have been increased by accumulation of yolk pools and tumours in the reproductive tract with age, which has previously been reported for *P. pacificus*, and which was found to be inhibited by mating with males (Kern et al. 2023). This is of interest as mating with males can also prolong the reproductive window and increase reproductive output in *C. elegans* (Pickett & Kornfeld 2013) and *P. pacificus* (C.W. personal observation), but whether this impacts the rupturing risk or its onset has not been explicitly examined.

Finally, it is also possible that the increased risk of rupturing seen in *P. pacificus* results from an age-dependent decline in osmoregulatory ability and/or cuticular integrity. The cuticle of *C. elegans* worms deforms and wrinkles with age (Herndon et al. 2002), and worms with aged or otherwise defective cuticles are at greater risk of rupturing under hypoosmotic stress (Rahimi et al. 2022, Chang et al. 2025). Testing these hypotheses in *Pristionchus* is beyond the scope of the present work, but it will be of particular interest to compare rupture-prone *P. pacificus* worms alongside rupture-resistant *P. fissidentatus* worms, as they produce a roughly similar number of progeny under the same osmotic and dietary conditions. The ultrastructure of the *P. fissidentatus* cuticle has not been examined in detail, but has been described as notably “thick” (Kanzaki et al. 2012), and this may bestow greater cuticular integrity that lowers rupturing risk, whatever the underlying cause of increased internal pressure.

Bagging (otherwise known as “matricidal hatching”, “bag-of-worms”, or “endotokia matricida”) also afflicts our two *Pristionchus* species differently, but in the opposite direction than seen in rupturing, being relatively common in *P. fissidentatus* and very rare in *P. pacificus*. Risk of bagging was highest, unsurprisingly, during the self-fertile reproductive window, while rupturing predominantly occurred as the self-fertile reproductive period was ending. As a result, we cannot rule out the possibility that the low rate of rupturing seen in *P. fissidentatus* is because worms that would have ruptured had already succumbed to bagging.

Work in *C. elegans* suggests that bagging is the result of senescent degeneration in vulval muscle function, and not obviously the result of damage caused by egg-laying, as progeny production was not predictive of whether bagging occurs (Pickett & Kornfeld 2013, Scharf et al. 2021). Because we housed our assay worms in small groups, not individually, we could not formally test for a role of progeny production rate/timing on bagging, but our cross-species findings argue against this hypothesis, given that the two *Pristionchus* species produce roughly similar numbers of progeny. As such, it seems more likely that the increased susceptibility to bagging seen in *P. fissidentatus* relative to *P. pacificus* reflects innate differences in age-dependent dysfunction of vulval musculature and/or sensory-feedback systems.

Placing fertile *C. elegans* hermaphrodites in starvation conditions can increase the rate of bagging (Chen & Caswell-Chen 2003), and this has been interpreted as an adaptive strategy of biomass sacrifice, with internally hatched eggs able to feed on maternal tissue and develop in periods of scarcity (Lohr et al. 2019). Whether this idea of adaptive sacrifice could drive variation among self-fertile *Pristionchus* species in bagging rate is hard to explain without knowing more about food availability and age-dependent mortality and reproductive rates under wild conditions; it is not established whether *Pristionchus* worms regularly reach the end of their self-fertile reproductive period in nature. However, the notion that bagging may be a part of an adaptive strategy, at least in some species, is supported by the recent discovery of a self-fertile *Pristionchus* species, *P. endotocus*, that bags “obligately” under laboratory conditions, and that is sometimes observed with a vulval plug that is hypothesised to hinder egg laying and help retain progeny inside the mother’s body (Güner et al. 2026). Despite this, extending this idea to dioecious species (which is most common, and represents the ancestral state for *Pristionchus*; Mayer et al. 2007) is challenging, as the obligate outcrossing and capacity for longer lifespans seen in these species (Weadick & Sommer 2016a) will combine to reduce the potential fitness benefits of sacrifice relative to those offered by surviving to mate and reproduce further.

Control of body size in nematodes is tightly associated with moulting, which they undergo four times as they mature, with rapid growth occurring during the periods when the protective but constraining cuticle is shed and rebuilt (Lažetić & Fay 2017). Growth also occurs between moults and after the final moult (Knight et al. 2002). Consistent with this, we found that *Pristionchus* hermaphrodites become significantly larger during adulthood, after the final moult, meaning body size can be a non-destructive indicator of worm age. Patterns of growth were slightly different between the two species: *P. pacificus* both lengthened and widened with age (by ∼25-30%), whereas the slightly larger *P. fissidentatus* grew wider by a similar degree without a clear increase in length, revealing divergent post-maturation growth trajectories within *Pristionchus*.

Body size in *C. elegans* is linked to growth of the epidermal syncytium, a giant multinucleate cell that surrounds the worm body and, alongside the seam cells, secretes the collagenous extracellular matrix that forms the cuticle (Lozano et al. 2006, Tuck 2014). While somatic cell divisions stop upon maturation in *C. elegans*, nuclear divisions continue within the epidermal syncytium. Endoreduplication can then cause further increases in cell ploidy and size, the extent of which can explain among-species variation in body size (Flemming et al. 2000). Age-specific patterns of collagen expression likely play a role in post-maturation growth dynamics as well. Mutant studies of cuticular collagens in *C. elegans* have shown contrasting effects on body size and shape, depending on the respective collagen’s structure and expression pattern. For instance, mutations in some lead to short and squat bodies (e.g., von Mende et al. 1988) whereas mutations in others result in long, thin bodies (e.g., Nyström et al. 2002). Furthermore, the expression of cuticular collagens changes with age in *C. elegans*; in general, collagen gene expression levels decline with age, but the pattern varies markedly across genes, with over an order of magnitude difference in the expression fold change between young and old adults (Wang et al. 2026). In light of these *C. elegans* findings, we speculate that the post-maturation growth observed in *P. pacificus* and *P. fissidentatus* is associated with endoreduplication of the epidermal syncytium, and the differences observed in body shape dynamics between the two reflects species-specific differences in age-dependent cuticular collagen expression patterns. The present study provides a foundation for testing these cellular and molecular hypotheses.

The nematode cuticle has functions beyond determining body size. It is a barrier that provides protection against environmental stressors, and it permits motility by establishing the worm’s hydrostatic skeleton (Johnstone 1994, Lints & Hall 2009, Murray et al. 2012). Given that, we tested for age effects on cuticular integrity and locomotory patterns. Cuticular integrity was tested by assaying whether permeability to Acridine Orange (AO) differed between young and old *P. pacificus* hermaphrodites. We observed a marked increase in the internalization of AO dye in older worms, indicating a weakening of the cuticle’s ultrastructural integrity. To rule out the possibility that this reflects increased ingestion, and not increased cuticular permeability, we also established that older *P. pacificus* worms, despite being bigger, have a lower feeding rate. This is consistent with previous findings of senescent cuticular integrity in *C. elegans* (Rahimi et al. 2022). As *C. elegans* ages, its cuticle thickens, wrinkles, and loses elasticity (Herndon et al. 2002, Rahimi et al. 2022), and these alterations appear to compromise cuticular strength. They also lead to increased susceptibility to environmental insults, such as bacteria, chemicals, and exogenous oxidative stress (Van de Walle et al. 2019). It should be noted that previous work found older *P. pacificus* worms were susceptible to cuticular blistering disease (Weadick & Sommer 2016a); however, this was not observed here, likely due to a different growth conditions in a new lab environment, including slightly different formulations of nematode growth media agar.

We also find locomotory performance is clearly age-linked in *P. pacificus* and *P. fissidentatus*, with crawling speed and turning rate decreasing with age, and wavelength increasing, in both species. This suggests that locomotory analysis provides a non-invasive approach to assaying ageing and overall health in *Pristionchus*. Our findings are consistent with past studies on *C. elegans*, which exhibits progressive decline in the ability to crawl normally with age (Herndon et al. 2002, 2017, Dhondt et al. 2021). An age-dependent decline in crawling speed was also seen in previous work on *P. pacificus*, an obligate outcrossing species that is a close relative of *P. pacificus*; notably, late-life crawling speed was associated with mating ability in older male *P. exspectatus* worms, but unrelated to mating ability in older females (Weadick & Sommer 2016b). Locomotion requires coordination of multiple physiological systems; altered locomotory behaviour could be due to senescent decline in any of these underlying systems. Given past work on *C. elegans*, the age-dependent changes seen in *Pristionchus* could be a result of a variety of age-associated changes: molecular damage resulting in injury or necrosis of the muscle tissue (Herndon et al. 2002, Glenn et al. 2004); age-dependent cuticle thickening, which reduces body flexibility (Bolanowski et al. 1981, Lints & Hall 2009, Page et al. 2014); or senescence in the underlying neurological system that drives crawling behaviour (Toth et al. 2012, Morsci et al. 2016).

Another possibility is that the altered locomotory behaviour of aged *Pristionchus* worms reflects a change in motivational state. Young adult *C. elegans* worms crawl faster, change direction more often, and have reduced postural wavelength when crawling on agar provisioned with food compared to bare agar (Angstman et al. 2016). These changes mirror the ones we observed for young versus old *Pristionchus* worms. Under this idea, older worms have less motivational drive to find and consume food, and they adjust their locomotory behaviour accordingly. Consistent with this idea, we found that feeding rate is diminished in older *P. pacificus* worms compared to younger ones, suggesting either decreased motivation or ability to feed. In *C. elegans*, age-dependent decline in feeding rate is associated with pathology of the pharynx and is a major contributor to mortality (Zhao et al. 2017). Interestingly, it is also known that reduced nutrient intake can trigger intestinal autophagy and promote longevity in *C. elegans* (Gelino et al. 2016), and this mechanism might explain the reduced intestinal integrity we observed in older versus young *Pristionchus* worms. Decreased ability to feed and/or absorb nutrients can be a suppressor of age-linked decline, depending on whether reduced calorie intake triggers lifespan-extending dietary restriction-related pathways (Bishop & Guarente 2007, Loo et al. 2023). Untangling causal mechanisms for senescence in *Pristionchus* will be a challenge, but the present study provides a baseline for future efforts.

## Conclusion

*Pristionchus* nematodes are an established model system for studying developmental genetics and evolution, and recent work suggests it may likewise prove useful for studying the biology of senescence. Here, we characterised several age-linked traits that can be used as healthspan indicators in *Pristionchus*, focusing primarily on well-studied *P. pacificus* but also its distantly related congener, *P. fissidentatus*. Our study reveals that ageing leads to large, frail-bodied nematodes with multiple dimensions of functional decline, and with varying degrees of consistency between the two species. This work will support further study into the molecular mechanisms of age-dependent decline in *Pristionchus* nematodes, and the evolutionary processes that shape patterns of senescence across species.

## Acknowledgements

This work was supported by a Royal Society Dorothy Hodgkin Fellowship Enhancement Award to CW.

## Bibliography

Angstman NB, Frank H-G, Schmitz C (2016) Advanced Behavioral Analyses Show that the Presence of Food Causes Subtle Changes in C. elegans Movement. Frontiers in Behavioral Neuroscience 10.

Arya U, Das CK, Subramaniam JR (2010) Caenorhabditis elegans for preclinical drug discovery. Current Science 99: 1669–1680.

Banse SA, Lucanic M, Sedore CA, Coleman-Hulbert AL, Plummer WT, Chen E et al. (2019) Automated lifespan determination across Caenorhabditis strains and species reveals assay-specific effects of chemical interventions. GeroScience 41: 945–960.

Bardgett RD, van der Putten WH (2014) Belowground biodiversity and ecosystem functioning. Nature 515: 505–511.

Bento G, Ogawa A, Sommer RJ (2010) Co-option of the hormone-signalling module dafachronic acid-DAF-12 in nematode evolution. Nature 466: 494–497.

Bishop NA, Guarente L (2007) Two neurons mediate diet-restriction-induced longevity in C. elegans. Nature 447: 545–549.

Blaxter M (2011) Nematodes: The Worm and Its Relatives. PLoS Biology 9: e1001050–e1001050.

Bolanowski MA, Russell RL, Jacobson LA (1981) Quantitative measures of aging in the nematode Caenorhabditis elegans. I. Population and longitudinal studies of two behavioral parameters. Mechanisms of Ageing and Development 15: 279–295.

Bulterijs S, Braeckman BP (2020) Phenotypic screening in C. Elegans as a tool for the discovery of new geroprotective drugs. Pharmaceuticals 13: 1–35.

Byerly L, Cassada RC, Russell RL (1976) The life cycle of the nematode Caenorhabditis elegans. Developmental Biology 51: 23–33.

Camargo A (2022) PCAtest: testing the statistical significance of Principal Component Analysis in R. PeerJ 10: e12967–e12967.

Chang H-W, Lin H-C, Yang C-T, Tay RJ, Chang D-M, Tung Y-C, Hsueh Y-P (2025) Cuticular collagens mediate cross-kingdom predator–prey interactions between trapping fungi and nematodes(AP Mitchell, Ed). PLOS Biology 23: e3003178.

Chen J, Caswell-Chen E (2003) Why Caenorhabditis elegans adults sacrifice their bodies to progeny. Nematology 5: 641–645.

Cinkornpumin JK, Hong RL (2011) RNAi mediated gene knockdown and transgenesis by microinjection in the necromenic nematode Pristionchus pacificus. Journal of Visualized Experiments.

Coburn C, Gems D (2013) The mysterious case of the C. elegans gut granule: death fluorescence, anthranilic acid and the kynurenine pathway. Frontiers in Genetics 4: 151–151.

Cox D (1972) Regression models and life tables (with discussion). J R Statist Soc B 34: 187–220-187–220.

Cox D, Oakes D (1984) Analysis of Survival Data, 1st eds. Chapman & Hall, New York.

Dhondt I, Verschuuren C, Zečić A, Loier T, Braeckman BP, De Vos WH (2021) Prediction of biological age by morphological staging of sarcopenia in Caenorhabditis elegans. Disease Models & Mechanisms 14.

Epstein J, Himmelhoch S, Gershon D (1972) Studies on ageing in nematodes III. Electronmicroscopical studies on age-associated cellular damage. Mechanisms of Ageing and Development 1: 245–255.

Félix MA, Hill RJ, Schwarz H, Sternberg PW, Sudhaus W, Sommer RJ (1999) Pristionchus pacificus, a nematode with only three juvenile stages, displays major heterochronic changes relative to Caenorhabditis elegans. Proceedings of the Royal Society B: Biological Sciences 266: 1617–1621.

Flemming AJ, Shen Z-Z, Cunha A, Emmons SW, Leroi AM (2000) Somatic polyploidization and cellular proliferation drive body size evolution in nematodes. Proceedings of the National Academy of Sciences 97: 5285–5290.

Friedman DB, Johnson TE (1988) A mutation in the age-1 gene in Caenorhabditis elegans lengthens life and reduces hermaphrodite fertility. Genetics 118: 75–86.

Fueser H, Mueller M-T, Traunspurger W (2020) Rapid ingestion and egestion of spherical microplastics by bacteria-feeding nematodes. Chemosphere 261: 128162–128162.

Gelino S, Chang JT, Kumsta C, She X, Davis A, Nguyen C, Panowski S, Hansen M (2016) Intestinal Autophagy Improves Healthspan and Longevity in C. elegans during Dietary Restriction. PLOS Genetics 12: e1006135–e1006135.

Gems D (2000) Longevity and ageing in parasitic and free-living nematodes. Biogerontology 1: 289–307.

Glenn CF, Chow DK, David L, Cooke CA, Gami MS, Iser WB, Hanselman KB, Goldberg IG, Wolkow CA (2004) Behavioral Deficits During Early Stages of Aging in Caenorhabditis elegans Result From Locomotory Deficits Possibly Linked to Muscle Frailty. The Journals of Gerontology Series A: Biological Sciences and Medical Sciences 59: 1251–1260.

Güner B, Kanzaki N, Weiler C, Rödelsperger C, Sumaya NHN, Sommer RJ, Herrmann M (2026) Description of Pristionchus endotocus n. sp., a new obligately bagging androdioecious species from the Philippines. Journal of Nematology 58: 78–93.

Harris JE, Crofton HD (1957) Structure and Function in the Nematodes: Internal Pressure and Cuticular Structure in Ascaris. Journal of Experimental Biology 34: 116–130.

Herndon LA, Altun ZF, Hall DH (2003) WormAtlas Glossary - B. WormAtlas.

Herndon LA, Schmeissner PJ, Dudaronek JM, Brown PA, Listner KM, Sakano Y, Paupard MC, Hall DH, Driscoll M (2002) Stochastic and genetic factors influence tissue-specific decline in ageing C. elegans. Nature 419: 808–814.

Herndon LA, Wolkow C, Driscoll M, Hall D (2017) Effects of Ageing on the Basic Biology and Anatomy of C. elegans. Ageing: lessons from C. elegans., 9–39. Springer International, Switzerland.

Hess KR (1995) Graphical methods for assessing violations of the proportional hazards assumption in cox regression. Statistics in Medicine 14: 1707–1723.

Hong RL, Sommer RJ (2006) Pristionchus pacificus: A well-rounded nematode. BioEssays 28: 651–659.

Hsin H, Kenyon C (1999) Signals from the reproductive system regulate the lifespan of C. elegans. Nature 399: 362–366.

Johnstone IL (1994) The cuticle of the nematode Caenorhabditis elegans: A complex collagen structure. BioEssays 16: 171–178.

Kanzaki N, Ragsdale EJ, Herrmann M, Sommer RJ (2012) Two New Species of Pristionchus (Rhabditida: Diplogastridae): P. fissidentatus n. sp. from Nepal and La Réunion Island and P. elegans n. sp. from Japan. Journal of nematology 44: 80–91.

Kaplan EL, Meier P (1958) Nonparametric Estimation from Incomplete Observations. Journal of the American Statistical Association 53: 457–481.

Kassambara A (2023) rstatix: Pipe-Friendly Framework for Basic Statistical Tests.

Kern CC, Srivastava S, Ezcurra M, Hsiung KC, Hui N, Townsend S et al. (2023) C. elegans ageing is accelerated by a self-destructive reproductive programme. Nature Communications 14: 4381.

Kern CC, Srivastava S, Ezcurra M, Hui N, Townsend S, Maczik D, Tse V, Bähler J, Gems D (2020) C. elegans hermaphrodites undergo semelparous reproductive death. bioRxiv: 2020.11.16.384255-2020.11.16.384255.

Kimura KD, Tissenbaum HA, Liu Y, Ruvkun G (1997) daf-2, an Insulin Receptor-Like Gene That Regulates Longevity and Diapause in Caenorhabditis elegans. Science 277: 942–946.

Knight CG, Patel MN, Azevedo RBR, Leroi AM (2002) A novel mode of ecdysozoan growth in *Caenorhabditis elegans*. Evolution & Development 4: 16–27.

Kumar S, Dietrich N, Kornfeld K (2016) Angiotensin Converting Enzyme (ACE) Inhibitor Extends Caenorhabditis elegans Life Span(SK Kim, Ed). PLOS Genetics 12: e1005866.

Lažetić V, Fay DS (2017) Molting in C. elegans. Worm 6: e1330246.

Lee DL (2002) The Biology of Nematodes, 1st ed. Taylor Francis, London.

Leiser SF, Jafari G, Primitivo M, Sutphin GL, Dong J, Leonard A, Fletcher M, Kaeberlein M (2016) Age-associated vulval integrity is an important marker of nematode healthspan. AGE 38: 419–431.

Lints R, Hall D (2009) The cuticle. WormAtlas.

Lionaki E, Tavernarakis N (2013) Assessing Aging and Senescent Decline in Caenorhabditis elegans: Cohort Survival Analysis. *Cell Senescence: Methods and Protocols*. Methods in Molecular Biology, 473–484. Springer Science+Business Media.

Lohr JN, Galimov ER, Gems D (2019) Does senescence promote fitness in Caenorhabditis elegans by causing death? Ageing Research Reviews 50: 58–71.

Loo J, Shah Bana MAF, Tan JK, Goon JA (2023) Effect of dietary restriction on health span in Caenorhabditis elegans: A systematic review. Experimental Gerontology 182: 112294–112294.

Lozano E, Sáez AG, Flemming AJ, Cunha A, Leroi AM (2006) Regulation of Growth by Ploidy in Caenorhabditis elegans. Current Biology 16: 493–498.

Luc M, Taylor DP, Netscher C (1979) On Endotokia Matricida and Intra-Uterine Development and Hatching in Nematodes. Nematologica 25: 268–274.

Mayer WE, Herrmann M, Sommer RJ (2007) Phylogeny of the nematode genus Pristionchus and implications for biodiversity, biogeography and the evolution of hermaphroditism. BMC Evolutionary Biology 7: 104–104.

von Mende N, Bird DM, Albert PS, Riddle DL (1988) dpy-13: A nematode collagen gene that affects body shape. Cell 55: 567–576.

Moczek AP, Sultan S, Foster S, Ledón-Rettig C, Dworkin I, Nijhout HF, Abouheif E, Pfennig DW (2011) The role of developmental plasticity in evolutionary innovation. Proceedings of the Royal Society B: Biological Sciences 278: 2705–2713.

Morgan K, McGaughran A, Villate L, Herrmann M, Witte H, Bartelmes G, Rochat J, Sommer RJ (2012) Multi locus analysis of Pristionchus pacificus on La Réunion Island reveals an evolutionary history shaped by multiple introductions, constrained dispersal events and rare out-crossing. Molecular Ecology 21.

Morsci NS, Hall DH, Driscoll M, Sheng Z-H (2016) Age-Related Phasic Patterns of Mitochondrial Maintenance in Adult Caenorhabditis elegans Neurons. The Journal of neuroscience : the official journal of the Society for Neuroscience 36: 1373–85.

Mosser T, Matic I, Leroy M (2011) Bacterium-Induced Internal Egg Hatching Frequency Is Predictive of Life Span in Caenorhabditis elegans Populations. Applied and Environmental Microbiology 77: 8189–8192.

Murray CJL, Vos T, Lozano R, Naghavi M, Flaxman AD, Michaud C et al. (2012) Disability-adjusted life years (DALYs) for 291 diseases and injuries in 21 regions, 1990–2010: a systematic analysis for the Global Burden of Disease Study 2010. The Lancet 380: 2197–2223.

Myers JA, Curtis BS, Curtis WR (2013) Improving accuracy of cell and chromophore concentration measurements using optical density. BMC Biophysics 6: 4.

Namai S, Sugimoto A (2018) Transgenesis by microparticle bombardment for live imaging of fluorescent proteins in Pristionchus pacificus germline and early embryos. Development Genes and Evolution 228: 75–82.

Namdeo S, Moreno E, Rödelsperger C, Baskaran P, Witte H, Sommer RJ (2018) Two independent sulfation processes regulate mouth-form plasticity in the nematode Pristionchus pacificus. Development 145: dev166272–dev166272.

Nyström J, Shen Z-Z, Aili M, Flemming AJ, Leroi A, Tuck S (2002) Increased or Decreased Levels of Caenorhabditis elegans lon-3, a Gene Encoding a Collagen, Cause Reciprocal Changes in Body Length. Genetics 161: 83–97.

Okkenhaug H, Chauve L, Masoudzadeh F, Okkenhaug L, Casanueva O (2020) Worm-align and Worm_CP, Two Open-Source Pipelines for Straightening and Quantification of Fluorescence Image Data Obtained from Caenorhabditis elegans. Journal of Visualized Experiments 159.

Page AP, Stepek G, Winter AD, Pertab D (2014) Enzymology of the nematode cuticle: A potential drug target? International Journal for Parasitology: Drugs and Drug Resistance 4: 133–141.

Park S-J, Goodman MB, Pruitt BL (2007) Analysis of nematode mechanics by piezoresistive displacement clamp. Proceedings of the National Academy of Sciences 104: 17376–17381.

Patel MN, Knight CG, Karageorgi C, Leroi AM (2002) Evolution of germ-line signals that regulate growth and aging in nematodes. Proceedings of the National Academy of Sciences 99: 769–774.

Photos A, Gutierrez A, Sommer RJ (2006) sem-4/spalt and egl-17/FGF have a conserved role in sex myoblast specification and migration in P. pacificus and C. elegans. Developmental Biology 293: 142–153.

Pickett CL, Kornfeld K (2013) Age-related degeneration of the egg-laying system promotes matricidal hatching in Caenorhabditis elegans. Aging Cell 12: 544–553.

Pires da Silva A (2006) Pristionchus pacificus genetic protocols. WormBook.

R Core Team (2019) R: a language and environment for statistical computing. R Foundation for Statistical Computing, Vienna, Austria.

Rae R, Iatsenko I, Witte H, Sommer RJ (2010) A subset of naturally isolated Bacillus strains show extreme virulence to the free-living nematodes Caenorhabditis elegans and Pristionchus pacificus. Environmental Microbiology 12: 3007–3021.

Rae R, Sinha A, Sommer RJ (2012) Genome-Wide Analysis of Germline Signaling Genes Regulating Longevity and Innate Immunity in the Nematode Pristionchus pacificus(JB Lok, Ed). PLoS Pathogens 8: e1002864–e1002864.

Rahimi M, Sohrabi S, Murphy CT (2022) Novel elasticity measurements reveal C. elegans cuticle stiffens with age and in a long-lived mutant. Biophysical Journal 121: 515–524.

Rödelsperger C, Neher RA, Weller AM, Eberhardt G, Witte H, Mayer WE, Dieterich C, Sommer RJ (2014) Characterization of Genetic Diversity in the Nematode *Pristionchus pacificus* from Population-Scale Resequencing Data. Genetics 196: 1153–1165.

Scharf A, Pohl F, Egan BM, Kocsisova Z, Kornfeld K (2021) Reproductive Aging in Caenorhabditis elegans: From Molecules to Ecology. Frontiers in Cell and Developmental Biology 9: 718522.

Schindelin J, Arganda-Carreras I, Frise E, Kaynig V, Longair M, Pietzsch T et al. (2012) Fiji: an open-source platform for biological-image analysis. Nature Methods 9: 676–682.

Schoenfeld D (1982) Partial residuals for the proportional hazards regression model. Biometrika 69: 239–241.

Serobyan V, Ragsdale EJ, Sommer RJ (2014) Adaptive value of a predatory mouth-form in a dimorphic nematode. Proceedings of the Royal Society B: Biological Sciences 281: 20141334–20141334.

Sieriebriennikov B, Prabh N, Dardiry M, Witte H, Röseler W, Kieninger MR, Rödelsperger C, Sommer RJ (2018) A Developmental Switch Generating Phenotypic Plasticity Is Part of a Conserved Multi-gene Locus. Cell Reports 23: 2835–2843.e4.

Simpson P (2002) Evolution of development in closely related species of flies and worms. Nature Reviews Genetics 3: 907–907.

Sinha A, Rae R, Iatsenko I, Sommer RJ (2012a) System wide analysis of the evolution of innate immunity in the nematode model species Caenorhabditis elegans and Pristionchus pacificus. PloS one 7: e44255–e44255.

Sinha A, Sommer RJ, Dieterich C (2012b) Divergent gene expression in the conserved dauer stage of the nematodes Pristionchus pacificus and Caenorhabditis elegans. BMC Genomics 13: 254–254.

Sommer RJ (2020) Phenotypic Plasticity: From Theory and Genetics to Current and Future Challenges. Genetics 215: 1–13.

Sommer RJ, Carta LK, Kim SY, Sternberg PW (1996) Morphological, genetic and molecular description of Pristionchus pacificus sp. n. (nematoda : neodiplogastridae). Fundamental and Applied Nematology 19: 511–521.

Stiernagle T (2006) Maintenance of C. elegans. WormBook.

Sun S, Rödelsperger C, Sommer RJ (2021) Single worm transcriptomics identifies a developmental core network of oscillating genes with deep conservation across nematodes. Genome Research 31: 1590–1601.

Sutphin GL, Backer G, Sheehan S, Bean S, Corban C, Liu T et al. (2017) *Caenorhabditis elegans* orthologs of human genes differentially expressed with age are enriched for determinants of longevity. Aging Cell 16: 672–682.

Tang Y, Horikoshi M, Li W (2016) ggfortify: Unified Interface to Visualize Statistical Results of Popular R Packages. The R Journal 8: 474–474.

Therneau T (2015) Survival: survival analysis. Version 2.38. https://CRAN.R-project.org/package=survival.

Toth ML, Melentijevic I, Shah L, Bhatia A, Lu K, Talwar A et al. (2012) Neurite sprouting and synapse deterioration in the aging Caenorhabditis elegans nervous system. The Journal of neuroscience : the official journal of the Society for Neuroscience 32: 8778–90.

Tuck S (2014) The control of cell growth and body size in Caenorhabditis elegans. Experimental Cell Research 321: 71–76.

Van de Walle P, Geens E, Baggerman G, José Naranjo-Galindo F, Askjaer P, Schoofs L, Temmerman L (2019) CEH-60/PBX regulates vitellogenesis and cuticle permeability through intestinal interaction with UNC-62/MEIS in Caenorhabditis elegans. PLOS Biology 17: e3000499–e3000499.

Vu V (2024) ggbiplot: A Grammar of Graphics Implementation of Biplots.

Wang AJ, Geppert BM, Beck D, Ji Y, Liu Y (2026) Collagen gene expression is linked to aging and lifespan extension in C. elegans. Matrix Biology Plus 30: 100192.

Weadick CJ, Sommer RJ (2016a) Mating system transitions drive life span evolution in Pristionchus nematodes. American Naturalist 187: 517–531.

Weadick CJ, Sommer RJ (2016b) Unexpected sex-specific post-reproductive lifespan in the free-living nematode Pristionchus exspectatus. Evolution and Development 18: 297–307.

Weadick CJ, Sommer RJ (2017) Hybrid crosses and the genetic basis of interspecific divergence in lifespan in Pristionchus nematodes. Journal of Evolutionary Biology 30: 650–657.

Weinkove D, Zavagno G (2021) Applying C. elegans to the Industrial Drug Discovery Process to Slow Aging. Frontiers in Aging 2.

Werner MS, Sieriebriennikov B, Loschko T, Namdeo S, Lenuzzi M, Dardiry M, Renahan T, Sharma DR, Sommer RJ (2017) Environmental influence on Pristionchus pacificus mouth form through different culture methods. Scientific Reports 7: 7207–7207.

Witte H, Moreno E, Rödelsperger C, Kim J, Kim JS, Streit A, Sommer RJ (2014) Gene inactivation using the CRISPR/Cas9 systemin the nematode Pristionchus pacificus. Development Genes and Evolution 225: 55–62.

Xiong H, Pears C, Woollard A (2017) An enhanced C. elegans based platform for toxicity assessment. Scientific Reports 7: 9839–9839.

Ye X, Linton JM, Schork NJ, Buck LB, Petrascheck M (2014) A pharmacological network for lifespan extension in C aenorhabditis elegans. Aging Cell 13: 206–215.

Zauner H, Mayer WE, Herrmann M, Weller A, Erwig M, Sommer RJ (2007) Distinct patterns of genetic variation in Pristionchus pacificus and Caenorhabditis elegans , two partially selfing nematodes with cosmopolitan distribution. Molecular Ecology 16: 1267–1280.

Zhao Y, Gilliat AF, Ziehm M, Turmaine M, Wang H, Ezcurra M et al. (2017) Two forms of death in ageing Caenorhabditis elegans. Nature Communications 8: 15458–15458.

