## Supplemental tables 1 and 2 for "Conserved and diverged patterns of senescence in *Pristionchus* nematodes"

Supplemental table 1: Summary of life history traits of *P. pacificus* and *P. fissidentatus*. Lifespan and age are in units of days since hatching.

| **Life history trait** | ***P. pacificus*** | ***P. fissidentatus*** |
|---|---|---|
| Median lifespan (95% CI) | 23 (23-25) | 21 (19-22) |
| Maximum lifespan | 44 | 32 |
| Individuals ruptured (%) | 29.0 | 12.8 |
| Individuals bagged (%) | 3.1 | 26.6 |
| Mean juveniles (± sd) | 164.5 ± 8.8 | 146.9 ± 11.7 |
| Maximum mean number of juveniles hatched on one day | 63.06 | 71.30 |
| The day of maximum number of juveniles | 5 | 5 |
| Age of start of reproduction | 3 | 3 |
| Latest age of reproduction | 9 | 8 |

Supplemental table 2: P values of locomotion metrics comparing young and older adults. ANOVA tests of locomotion metric data comparing worm age within each species. A Levene's Test for Homogeneity of Variance revealed track length of older adult *P. pacificus* is more variable than younger adults ($F(1, 34) = 11.683$, $p = 0.001$), as is amplitude in *P. fissidentatus* ($F(1, 53) = 6.260$, $p = 0.014$). *** = $p < 0.001$, ** = $p < 0.01$, * = $p < 0.05$. All df1 = 1. ns = not significantly different. In *P. pacificus* df2 = 34, and *P. fissidentatus* df2 = 53.

| Species and metric | Young adult mean ± sd | Old value mean ± sd | F | P | Mean change |
|---|---|---|---|---|---|
| *P. pacificus* | | | | | |
| Track length (µm) | 3749 ± 780.0 | 3801 ± < 780.0 | 0.038 | 0.847 | ns |
| Wavelength (µm) | 497± 51.8 | 568 ± 70.5 | 17.585 | <0.001 *** | + 70.68 |
| Amplitude (µm) | 64.3 ± 21.8 | 71.3 ± 21.6 | 2.069 | 0.157 | ns |
| Turn count (turns / min) | 34 ± 15.5 | 22 ± 12.2 | 10.270 | 0.002 ** | - 12 |
| Mean speed (µm/s) | 130.63 ± 14.8 | 92.52 ± 21.50 | 12.770 | <0.001 *** | - 38.11 |
| Maximum speed (µm/s) | 723 ± 292 | 552 ± 288 | 6.028 | 0.017 * | - 171 |
| *P. fissidentatus* | | | | | |
| Track length (µm) | 2464 ± 645.8 | 3710 ± 941.3 | 37.522 | <0.001 *** | + 1246 |
| Wavelength (µm) | 577 ± 85.7 | 636 ± 78.1 | 11.632 | <0.001 *** | + 59.13 |
| Amplitude (µm) | 73.9 ± 27.2 | 82.7 ± 39.4 | 0.901 | 0.345 | ns |
| Turn count (turns / min) | 17 ± 8.40 | 11 ± 4.5 | 18.756 | <0.001 *** | - 6 |
| Mean speed (µm/s) | 107.62 ± 11.3 | 81.53 ± 11.32 | 20.310 | <0.001 *** | - 26.09 |
| Maximum speed (µm/s) | 605 ± 338 | 451 ± 294 | 6.628 | 0.011 * | - 154 |
